# Counteracting Genome Instability by p53-dependent Mintosis

**DOI:** 10.1101/2020.01.16.908954

**Authors:** Jianqing Liang, Zubiao Niu, Xiaochen Yu, Bo Zhang, You Zheng, Manna Wang, Banzhan Ruan, Hongquan Qin, Xin Zhang, Songzhi Gu, Xiaoyong Sai, Yanhong Tai, Lihua Gao, Li Ma, Zhaolie Chen, Hongyan Huang, Xiaoning Wang, Qiang Sun

**Author notes:** Correspondence to: Qiang Sun, Xiaoning Wang, Hongyan Huang. These authors contributed equally to this work (Jianqing Liang, Zubiao Niu, Xiaochen Yu).

## Abstract

Entosis was proposed to promote aneuploidy and genome instability by cell-in-cell mediated engulfment in tumor cells. We reported here, in epithelial cells, that entosis coupled with mitotic arrest functions to counteract genome instability by targeting aneuploid mitotic progenies for engulfment and elimination. We found that the formation of cell-in-cell structures associated with prolonged mitosis, which was sufficient to induce entosis. This process was controlled by the tumor suppressor p53 (wild type) that upregulates Rnd3 expression in response to DNA damages associated with prolonged metaphase. Rnd3 compartmentalized RhoA activities accumulated during prolonged metaphase to drive cell-in-cell formation. Remarkably, this prolonged <u>m</u>itosis-<u>in</u>duced en<u>tosis</u> (mintosis) selectively targets non-diploid progenies for internalization, blockade of which increased aneuploidy. Thus, our work uncovered a heretofore unrecognized mechanism of mitotic surveillance for entosis, which eliminates newly-born abnormal daughter cells in a p53-depedent way to maintain genome integrity.

## Introduction

Cell division via mitosis is strictly regulated to ensure the production of healthy daughter cells. During mitosis, correct segregation and distribution of genetic materials into daughter cells rely on proper activation of spindle assemble checkpoint (SAC) (Musacchio, 2015; Sivakumar & Gorbsky, 2015). SAC is activated by unsatisfied microtubule-kinetochore attachment, leading to the recruitment of SAC proteins MAD2, BUBR1 and BUB3 together with CDC20 (Kapanidou *et al*, 2017) to kinetochores to form mitotic checkpoint complex (MCC). MCC functions to inhibit anaphase-promoting complex or cyclosome (APC/C), the multisubunit E3 ubiquitin ligase essential for metaphase-anaphase transition in the presence of CDC20. Molecules disrupting MCC such as CUEDC2 could activate APC/C via preventing MAD2 from binding to CDC20 (Gao *et al*, 2011). Activated APC/C promotes proteasome-dependent degradation of cyclin B1 (CCNB1) to allow mitotic exit, and securin to release separase (ESPL1) which leads to cohesion destruction and subsequent sister chromatid separation (Musacchio, 2015; Sivakumar & Gorbsky, 2015). Prolonged SAC activation due to mitotic aberrations generally leads to mitotic arrest and often, but not always, to mitotic catastrophe (specifically referred to catastrophic death here) (Vitale *et al*, 2011). Those that eventually pass mitotic arrest are prone to produce aneuploid/polyploid (non-diploid) daughter cells that need to be dealt with post-mitotically, otherwise impair tissue homeostasis and/or contribute to tumorigenesis (Funk *et al*, 2016; Santaguida & Amon, 2015).

Entosis is a recently defined non-apoptotic cell death program, where suspended epithelial cells actively penetrate into and die inside of their neighbors (Overholtzer *et al*, 2007). Cell penetration leads to the formation of so called “cell-in-cell” structures (CIC), which requires polarized actomyosin contraction at the rear cortex of the internalizing cells. The polarized distribution of actomyosin is established by local inhibition of RhoA-ROCK signaling at cell-cell junctions, where lie junction-associated inhibitors, such as p190A RhoGAP (Sun *et al*, 2014a). Multiple factors, including CDKN2A (Liang *et al*, 2018), PCDH7 (Wang *et al*, 2020), IL-8 (Ruan *et al*, 2018a) and membrane cholesterol and lipids (Ruan *et al*, 2018b), that affects RhoA signaling were recently identified as important regulators of entotic CIC formation. Although initially viable, majority of the internalized cells die non-autonomously with the assistance of outer host cells (Florey *et al*, 2011). Pathologically in the context of tumors, entosis was proposed as a cellular mechanism of cell competition that selects winner tumor cell clones via internalizing and killing loser cells (Kroemer & Perfettini, 2014; Sun *et al*, 2014b), and promotes tumor evolution by inducing genome instability of outer cells (Krajcovic *et al*, 2011; Mackay *et al*, 2018). However, its roles in physiological context remain largely speculative and mysterious up to date.

The tumor suppressor p53 was known as “guardian of genome”. In responding to acute DNA damages, signaling cascades involving ATM/ATR and/or CHK1/CHK2 were initiated to activate p53, which was believed to triggers cellular responses such as cell cycle arrest or apoptosis to counteract the genotoxic stresses (Bieging *et al*, 2014). p53 initiates different cellular responses generally via regulating expression of specific downstream target genes, such as the CDK inhibitor *p21* for cell cycle arrest and the pro-apoptotic Bcl-2 family members *Puma* and *Noxa* for apoptosis (Joerger & Fersht, 2016). While cell cycle arrest was envisaged to provide cells an opportunity to fix the repairable DNA damages before next round of cell cycle and thus prevent propagation of potentially harmful mutations, cell death by apoptosis was likely a more aggressive and efficient way to eliminate cells harboring more severe genetic aberrations, and thereby to maintain genome integrity (Bieging *et al.*, 2014; Joerger & Fersht, 2016). In addition to apoptosis, p53 was also implicated in other forms of cell deaths that play roles in removing cells with DNA damages (Kruiswijk *et al*, 2015). Mitotic catastrophe was believed to be a p53-regulated non-apoptotic cell death that eliminates questionable cells during prolonged mitosis which generally takes place with DNA damages (Ranjan & Iwakuma, 2016). Nevertheless, it’s unknown whether genetically questionable progenies following mitosis were actively removed by a non-apoptotic cell death that might also be regulated by p53.

Here, we demonstrate that p53 mediates the internalization and elimination of aneuploidy daughter cells from transient mitotic arrest by a non-apoptotic death program termed mintosis (derived from <u>p</u>rolonged <u>m</u>itosis-<u>in</u>duced en<u>tosis)</u>, which was driven by Rnd3-compartmentalized RhoA activities in daughter cells that were internalized in response to DNA damages during prolonged metaphase. Thus, our work identified a novel surveillance mechanism for entosis, in a wild type p53-depedent way, to safeguard mitosis and genome integrity.

## Results

### Entosis is Associated with Prolonged Mitosis

In a process of tracking entotic CIC formation by time lapse microscopy in adherent MCF10A, a non-transformed human mammary epithelial cell line routinely used for entosis research, we unexpectedly found that majority (22/24) of CIC structures formed shortly after mitotic cell division, in spite of a few of exceptions where one adherent cell migrating into and one suspended cells sinking into their neighbors (Figure 1A, C and Movie S1). Once internalized, the inner cells readily underwent cell death (Figure 1B, D, Figure S1 and Movie S2-3) as previously reported for entosis (Overholtzer *et al.*, 2007). The result suggests that mitosis could initiate entosis in adherent monolayers as recently reported for the induced entosis, via CDC42 depletion, in 16HBE cells (Durgan *et al*, 2017). However, only a small portion of the mitotic events (<0.5% or so) led to entosis in normal adherent MCF10A culture, suggesting that entotic mitosis, referring to mitosis leading to entosis, is intrinsically different from normal mitosis. Comparison analysis revealed a significantly prolonged metaphase in entotic mitosis of MCF10A cells (Figure 1A, 1E and Movie S1, 2), Similar entosis also occurred in MCF7, a breast cancer cell line that frequently undergoes entosis (Figure S1C, S1D), indicating that entosis may take place in respond to mitotic arrest.

**Figure 1.**
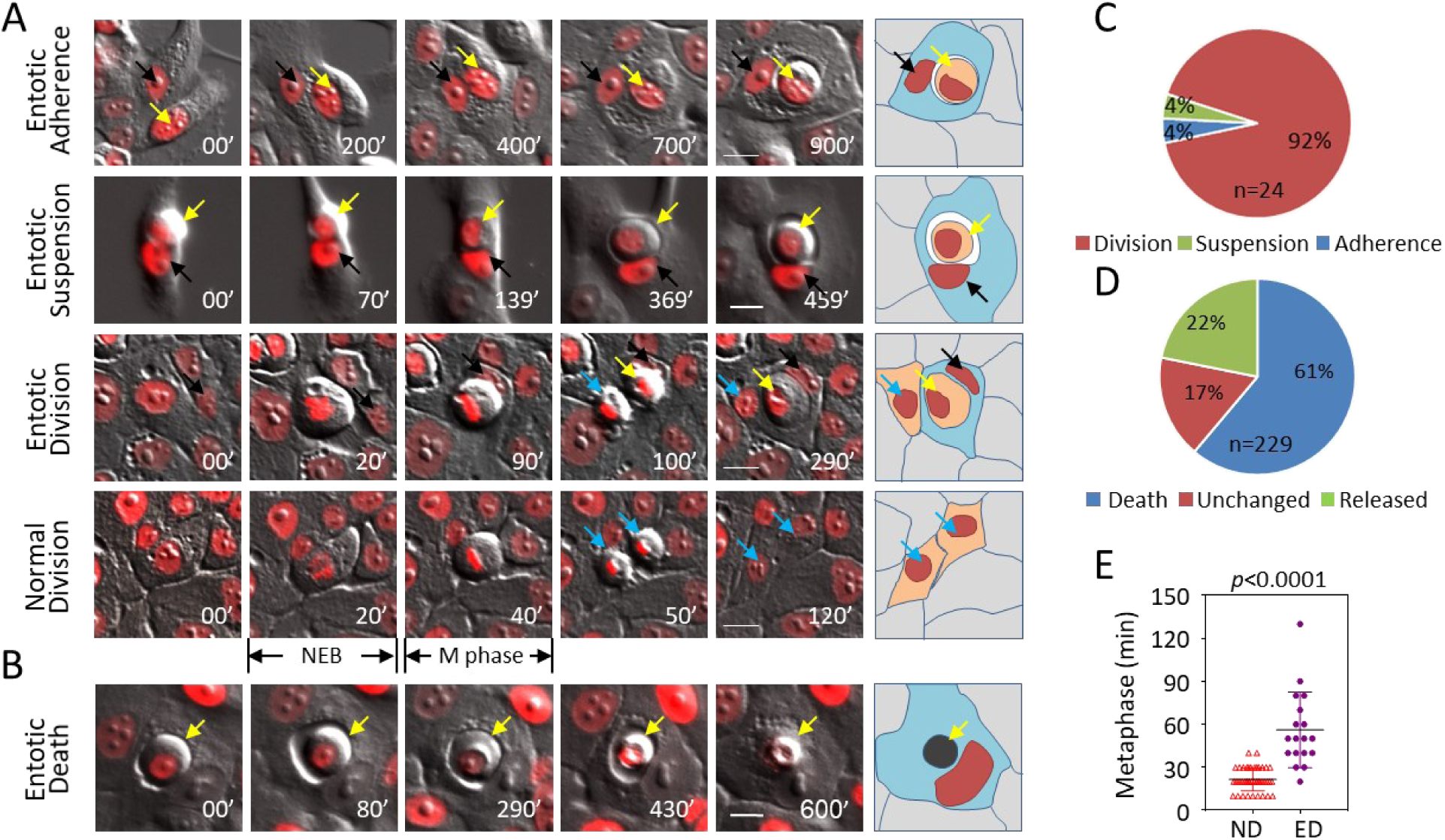
Entosis is preceded by mitosis of prolonged metaphase. (A) Representative image sequences for the formation of entotic CIC structures from adherent cell (panel of Entotic Adherence), suspended cells (panel of Entotic Suspension), divided cells (panel of Entotic Division), and normal cell division (panel of Normal Division). Yellow arrows indicate internalized cells, black arrows indicate outer cells, blue arrows indicate cells adhered to plate bottom. NEB: nuclear envelop breakdown; Scale bar: 20 μm. Also see Movie S1. (B) Representative image sequences for inner cell death of entotic CIC structures in MCF10A cells. Arrows indicate inner cell. Scale bar: 20 μm. (C) Quantification of Entotic Adherence (Adherence), Entotic suspension (Suspension) and Entotic Division (Division). Quantification of inner cell fates in entotic CIC structures over 24 h period in MCF10A cells. Also see Movie S3. (D) Metaphase analysis of normal cell division (ND) and entotic cell division (ED) referring to cell division leading to entotic CIC formation in MCF10A cells. n=52 for ND, 18 for ED.

### Transient Mitotic Arrest Activates Entotic CIC Formation

To test this idea, we performed RNAi-mediated knockdown (KD) of CDC20 (Figure 2A) and ESPL1 (Figure S2A), two core genes critical for metaphase to anaphase transition, in MCF10A cells. Both CDC20 and ESPL1 depletion efficiently induced mitotic arrest of various extents (Figure 2B and S2B-C) and importantly increased CIC formation 36 hours post siRNA transfection (Figure 2C and S2D) when a number of mitotically arrested cells started to appear. Similar to that in spontaneous entosis, mitosis is corresponding to majority of CIC formation (47/51 for CDC20 KD, 53/54 for ESPL1 KD) for these two induced entosis (Figure 2D and S2E), consistent with which, blocking cell cycle with CDKs’ inhibitors (Ro-3306 and SU-9516) efficiently inhibited CIC formation (Figure 2C). Interestingly, while metaphase of normal division peaks around 20 min for MCF10A cells (Figure 2B, 2E and S2C), entotic mitosis displays a restricted metaphase range from 30 min to 195 min (Figure 2E and S2C) with most entotic events preceded by mitotic arrest of 60 min or so. Mitotic cells arrested for more than 200 min were unlikely capable of division and eventually ended up with catastrophic death (Figure 2E, 2F, S2C and Movie S4). Therefore, mitotic catastrophe and entosis are likely cooperated to safeguard aberrant mitosis with each of them worked before and after mitosis, respectively. To further confirm the role of mitotic arrest in activating entosis, mitotic arrest induced by depleting CUEDC2, a promoter of metaphase-anaphase transition (Gao *et al.*, 2011), was released by co-depleting either BUBR1 or MAD2 (Figure 2G and S2F-H), two essential components of mitotic checkpoint complex (Sivakumar & Gorbsky, 2015). As a result, mitosis-activated CIC formation induced by CUEDC2 depletion was inhibited as well (Figure 2H-I). The impact of mitotic arrest on entosis induction was also confirmed in MCF7 cells (Figure S2I, S2J). Together, the data presented above supports that mitotic arrest, indicated by prolonged metaphase, primes cells to undergo entotic CIC formation post-mitotically. To differentiate from entosis induced via other ways, we refer to this prolonged <u>m</u>itosis-<u>in</u>duced en<u>tosis</u> as mintosis hereafter.

**Figure 2.**
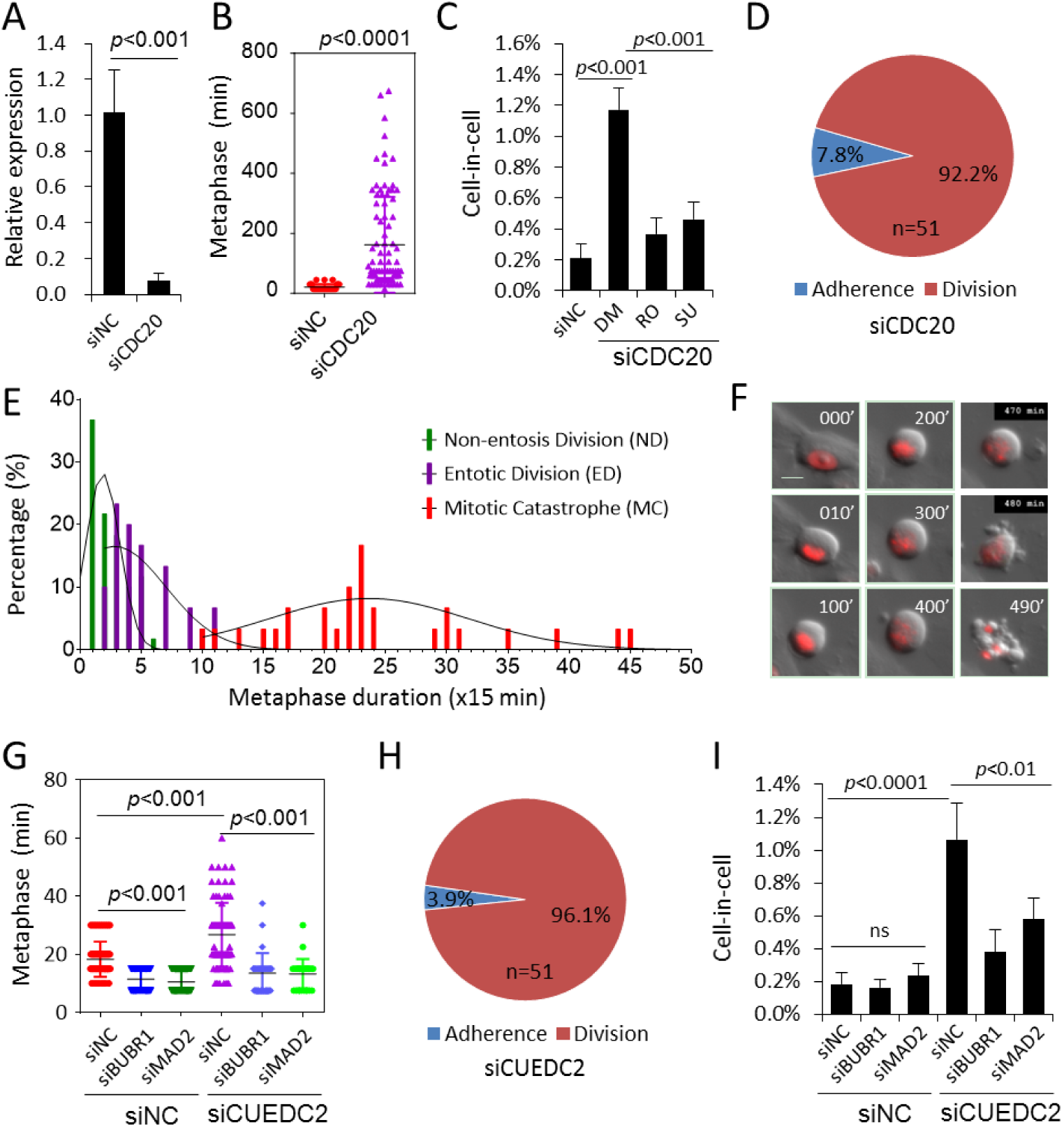
Prolonged mitosis primes cells to undergo entosis. (A) CDC20 mRNA level examined by quantitative PCR (qPCR) upon knockdown by RNA interference. Data are mean ± SD of triplicate experiments. *p<*0.001. (B) Graph plots metaphase duration of control (siNC, n=30) and CDC20 depleted (siCDC20, n=90) cells. *p<*0.0001. (C) Quantification of CIC structures in control (siNC) and CDC20 (siCDC20) depleted cells. DM: DMSO; RO: Ro-3306(10 μM); SU: SU-9516 (5 μM. Data are mean ± SD of 4 or more fields with more than 5000 cells analyzed each. *p<*0.001 for the pairs of siNC vs DM, DM vs RO, DM vs SU. (D) Quantification of Entotic Adherence (Adherence) and Entotic Division (Division) in MCF10A cells with CDC20 depletion. (E) Histogram analysis with fitted curves of metaphase from non-entosis division (ND), entotic division (ED) and mitotic catasphrohpe (MC).n=30 for ND and MC, 24 for ED. Gaussian curve was created by nonlinear regression of the frequency distribution by GraphPad Prism software. (F) Image sequence shows example of mitotic catastrophe with 490 min metaphase. Also see Movie S4. Scale bar: 20 μm (G) Graph plots metaphase duration of control (siNC) and CUEDC2, BUBR1 and MAD2 depleted cells. From left to right, n=58, 60, 60, 85, 30, 30, respectively. *p<*0.001 between pairs analyzed as indicated. (H) Quantification of Entotic Adherence (Adherence) and Entotic Division (Division) in MCF10A cells with CUEDC2 depletion. (I) Quantification of CIC structures in cells co-depleting CUEDC2 with BUBR1 or MAD2 genes. Data are mean ± SD of 4 or more fields with more than 5000 cells analyzed each. ns: not significant. *p<*0.0001 between siCUEDC2 and siNCs, *p<*0.01 between siNC and siBUBR1 or siMAD2 within siCUEDC2 group.

### DNA Damages Promote Mintosis

DNA damages were reported being associated with prolonged mitosis (Ganem & Pellman, 2012). We therefore hypothesized that DNA damages might account for mintosis incurred by prolonged mitosis. Consistent with this idea, time lapse-associated immunostaining indicated that cells of longer metaphase were positive in γH2AX (Figure 3A-B, Figure S3A), a marker of DNA damages, with nuclear γH2AX foci significantly more than those in cells of shorter metaphase (Figure 3C). Moreover, the nuclear γH2AX foci were correlated with formation of mintotic CIC structures, where the internalizing mitotic daughter cells contained more nuclear γH2AX foci than their siblings that didn’t form CIC structures (Figure 3D). Significantly, mintotic CIC formation was consistently suppressed by γH2AX depletion (Figure 3E-F) or inhibition of its downstream signaling by chemical compounds (Figure 3G) or siRNAs targeting ATM, ATR, CHK1 and CHK2 (Figure S3B-C), indicating an essential role of DNA damage signaling in mintosis. To directly examine the effects of DNA damages on mintosis, cells synchronized in M phase were treated with mitomycin, a DNA damage inducer, and then cultured in normal media to allow mitotic exit and cytokinesis. As shown Figure 3H-I, mitomycin treatment efficiently activated DNA damage response, indicated by enhanced expression of γH2AX and phosphorylated ATM (Figure 3H), and significantly increased mintotic CIC formation following cell division (Figure 3I). These effects were confirmed by bleomycin, another DNA damage inducer (data not shown). Interestingly, DNA damage signaling was likely also activated in human breast cancer tissues with high CIC structures, where nuclear γH2AX foci were significantly more than those in low-CIC breast cancer tissues (Figure 3J-K). Together, these data suggest that prolonged mitosis-associated DNA damages induce mintosis.

**Figure 3.**
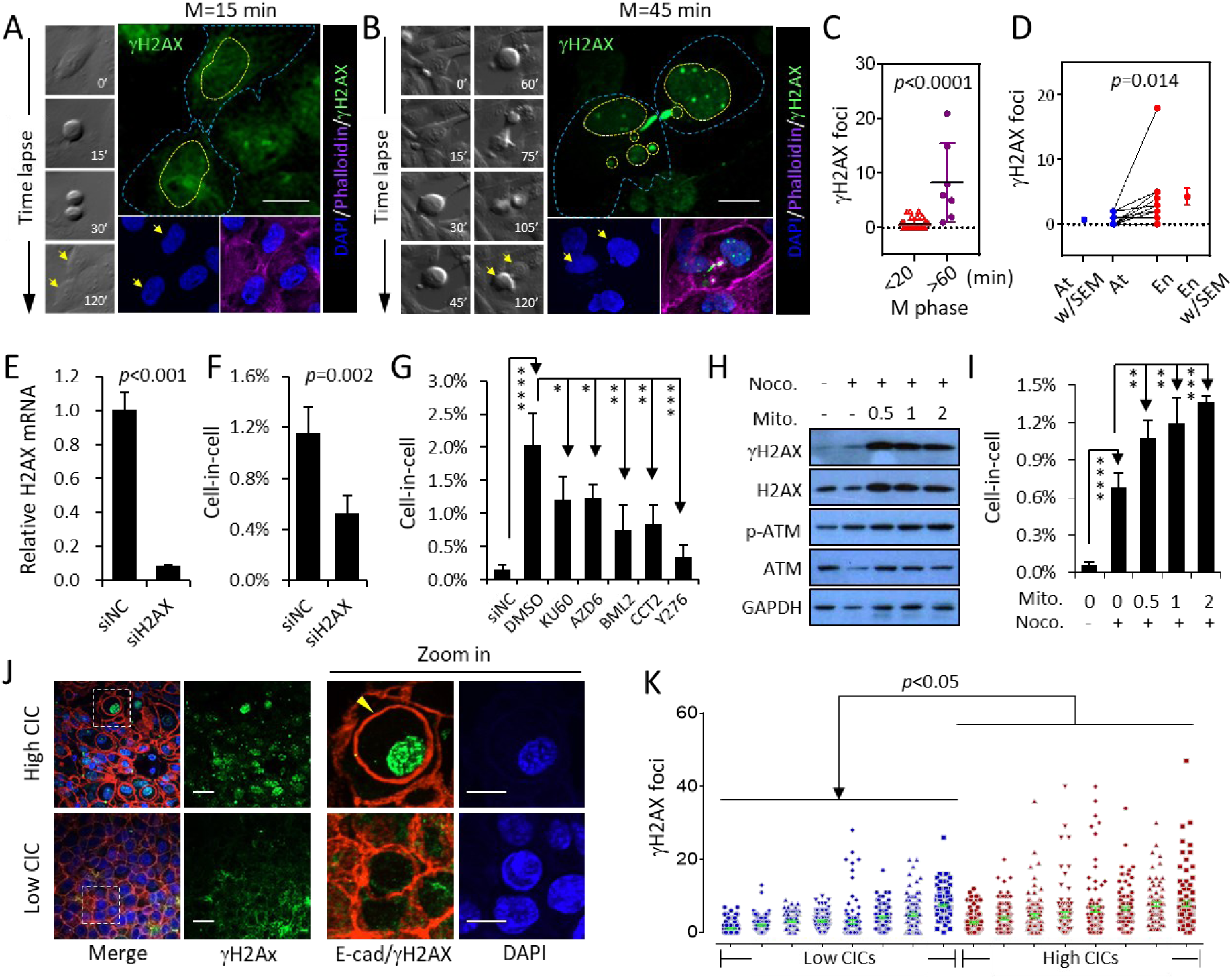
Mitotic DNA damages promote mintosis. (A-B) Representative images for time lapse-associated immunostaining of γH2AX in mitotic cells of short M phase (A, 15 min) and long M phase (B, 45 min), with corresponding DIC images of time lapse on the left and images for DAPI and merge channels on the bottom. Arrows indicate sibling cells. Blue dashed lines depict cell shape, yellow dashed lines depict shape of nuclei. Note: sibling cells from long M phase (B, 45 min) contain micronuclei indicating chromosome missegregation, and multiple γH2AX foci indicating DNA damages. Scale bar: 10 μm. (C) Number of nuclear γH2AX foci in cells from mitosis of different M phases. n=48 for M phase<20 min and 7 for M phase>60 min. (D) Number of nuclear γH2AX foci in paired sibling cells with one attached to the culture bottom (At) while another internalized to form CIC structure (En). n=12. (E-F) H2AX depletion (E) inhibits mintotic CIC formation (F). Data are mean ± SD of 4 or more fields with more than 5000 cells analyzed each. (G) Effects of inhibiting DDR signaling on mintotic CIC formation in CUEDC2-depleted cells by compounds targeting ATM (KU60 for KU-60019), ATR (AZD6 for AZD6738), CHK1 (BML2 for BML-277) and CHK2 (CCT2 for CCT245737). Inhibition of ROCKs by Y27632 (Y276) as positive control. Data are mean ± SD of 4 or more fields with more than 4000 cells analyzed each. siNC for non-target control siRNA. (H) DNA damages by mitomycin (Mito.) increase expression of γH2AX and phospho-ATM in cells synchronized in M phase by nocodazole (Noco.). (I) Formation of mintotic CIC structures in cells 18 hours released from mitotic arrest under conditions in (H). Data are mean ± SD of 4 or more fields with more than 4000 cells analyzed each. (J) Representative images for γH2AX staining in breast cancer tissues. E-cadherin (E-cad) in red indicates cell junctions. Arrow head indicates inner cell of a CIC structure. Scale bars: 20μm for the left, 10μm for zoomed images in the right. (K) Quantification of nuclear γH2AX foci in 16 human breast cancer samples. About one hundred of cells were quantified for each sample.

### P53 is Required for Daughter Cells to Undergo Mintosis

Since p53 is the key mediator of DNA damage signaling, we then explore the involvement of p53 in mintosis. Time lapse-associated immunostaining indicated that high level of p53 accumulated in the nuclei of daughter cells penetrating into their neighbors (Figure 4A). The expression levels of p53 were positively correlated with γH2AX foci formation (Figure 4B), and higher in mitotic cells of longer metaphase (Figure 4C), and higher in the internalizing daughter cells as compared with their non-mintotic sibling cells (Figure 4D) and those from normal division (Figure 4E). Importantly, RNA interference-mediated knockdown of p53 significantly suppressed mintotic CIC formation induced by CDC20 or CUEDC2 knockdown (Figure 4F-G), suggesting that p53 is required for mintosis. By a coculture experiment, p53 was demonstrated to function primarily in the mitotic daughter cells that penetrate into their neighbors, as p53 knockdown reduced the frequency of cells’ penetration as inners when co-cultured with control cells (Figure 4H-J).

**Figure 4:**
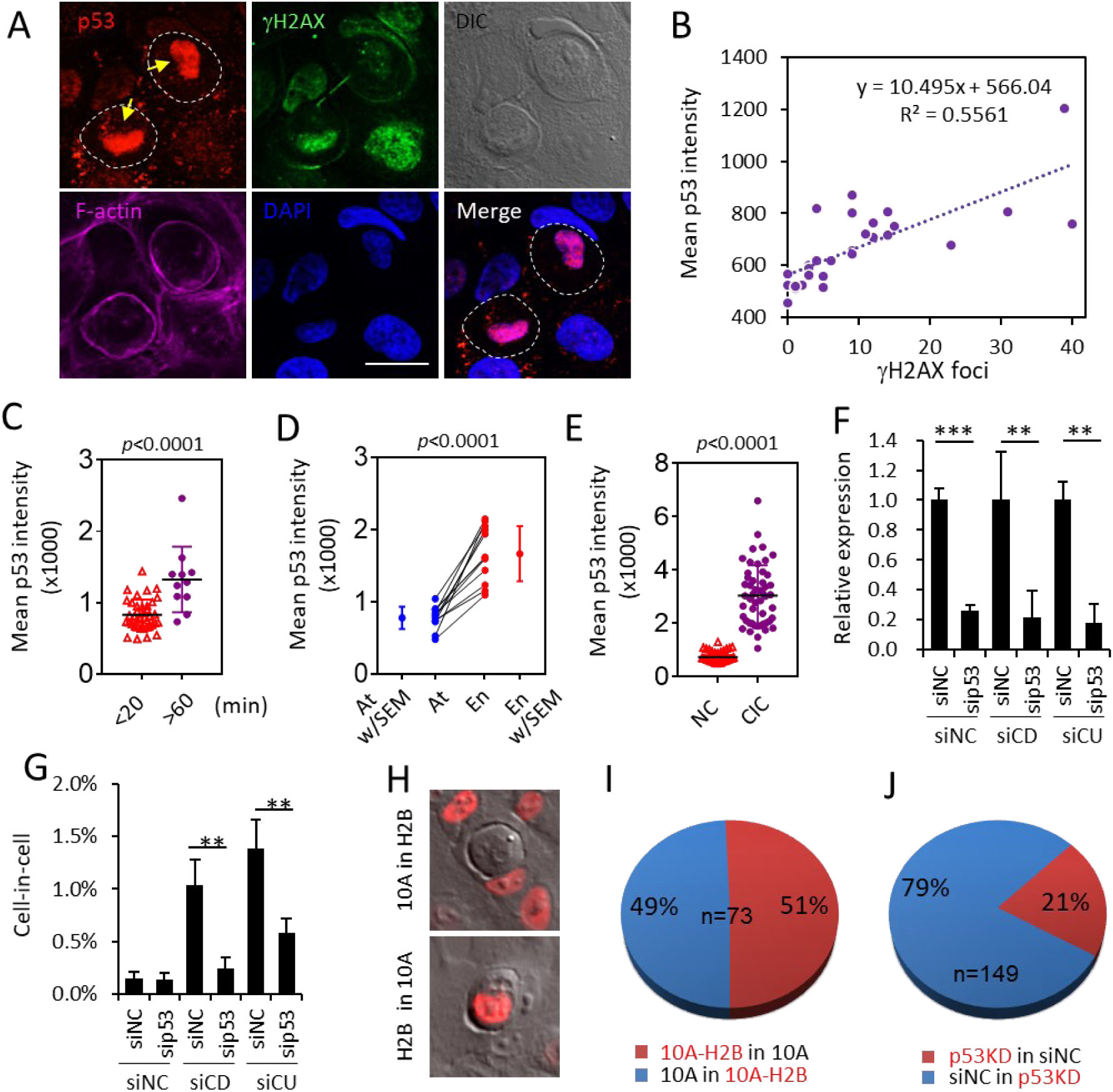
p53 signaling pathway is required for mintosis. (A) Representative images for p53 staining in daughter cells forming CIC structures as indicated by yellow arrows. Scale bar is 10μm. (B) Correlation analysis between p53 expression and number of nuclear γH2AX foci in MCF10A cells. n=30. (C) Expression of p53 as indicated by mean staining intensity in cells from mitosis of different M phases. n=38 for M phase<20 min and 11 for M phase>60 min. (D) (D) Expression of p53 in paired sibling cells with one attached to the culture bottom (At) while another internalized to form CIC structure (En). n=12. (E) Expression of p53 in inner cells of fresh CIC structures (CIC) is higher than that in single mono-nucleic cell (NC). n=50 for each group. (F-G) p53 depletion (sip53) inhibits mintotic CIC formation induced by CDC20 (siCD) and CUEDC2 (siCU) knockdown. siNC for non-target control siRNA. (H-J) Reduced frequency for p53 knockdown (p53KD) cells (labeled with H2B-mCherry in (J)) to penetrate into control cells (siNC). siNC for non-target control siRNA.

### RhoA Activity Accumulated with Prolonged Metaphase Promotes Mintosis

To explore the potential mechanisms underlying the regulation of mintosis by p53, we first examine changes in RhoA activities, which is prerequisite for CIC formation (Ning *et al*, 2015; Sun *et al.*, 2014a) and regulated by p53 signaling (Gadea *et al*, 2007; Xia & Land, 2007), during mintosis. CDC20-depleted cells were stained with antibody for p-MLC, a readout of RhoA activities, following 4-hour time lapse microscopy (Movie S5). As shown in Figure 5A-C, mitotic cells of longer metaphase tended to display higher p-MLC intensity, in agreement with which, RhoA activities in mitotic cells did accumulate over time as monitored by FRET time lapse analysis (Figure 5D, S4A-B and Movie S6). Since mintotic CIC formation occurred in daughter cells, we examined whether enhanced RhoA activities in mother cells could be inherited by daughter cells. RhoA activities of daughter cells right after cytokinesis were compared with those of their respective mother cells in the end of metaphase by FRET time lapse. As shown in Figure S4A-D, the two sibling daughter cells are tightly adherent to their mothers in RhoA activities, which is true for both normal and mintotic cell division, suggesting that RhoA activities could be transmitted from mother cells to daughters with little loss. We then examined the effects of changed RhoA activities on mintotic CIC formation, consistent with essential role of RhoA in entosis, ectopic overexpression of RhoA significantly increased mintotic CIC frequency (Figure 5E), and inhibition of RhoA signaling by Y27632 targeting ROCKs efficiently suppressed mintotic CIC formation (Figure 5F), supporting a role of increased RhoA activity in promoting mintosis.

**Figure 5.**
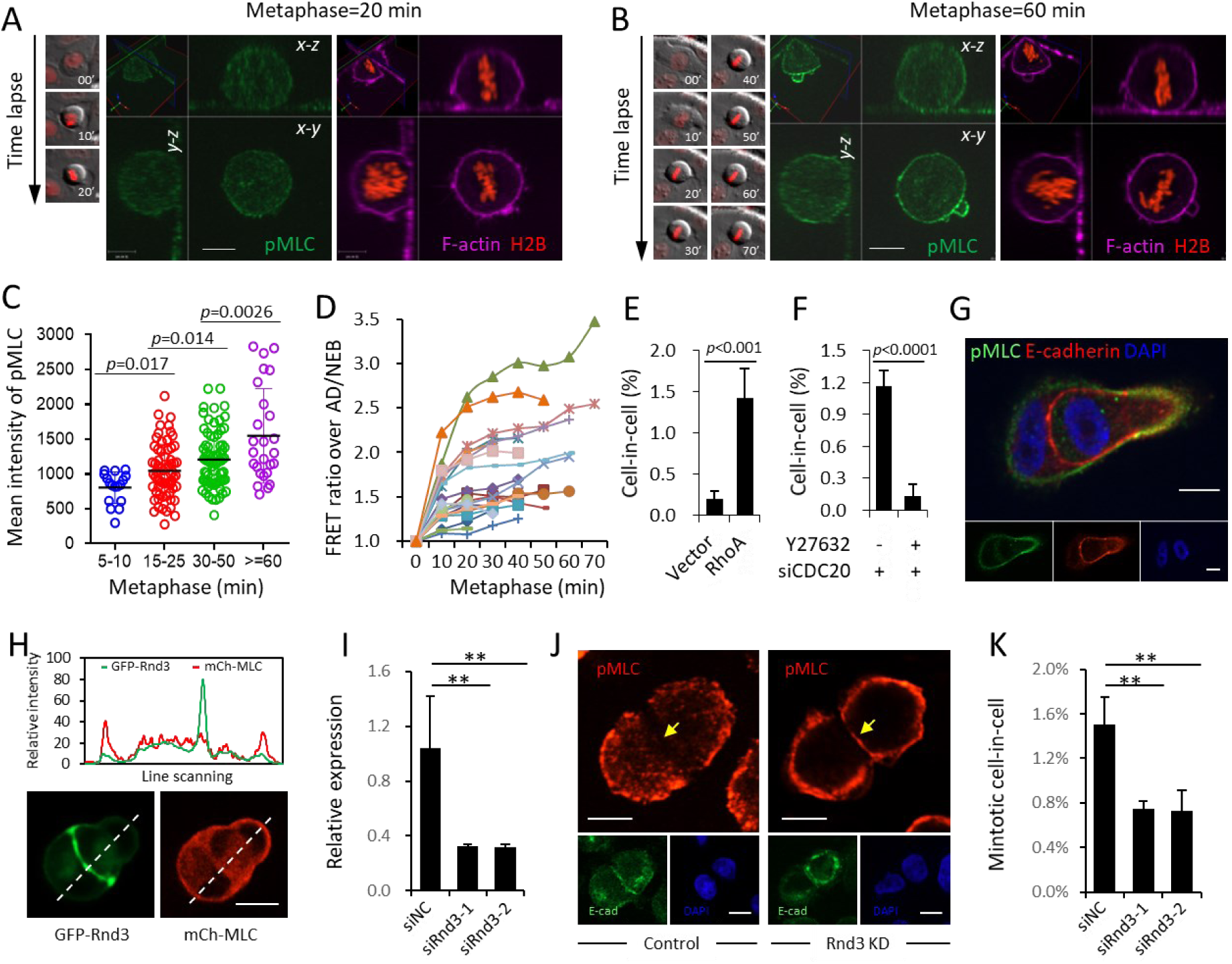
Rnd3 is required for asymmetric RhoA activation and mintotic CIC formation. (A-B) Representative image of pMLC staining in mitotic cell of normal metaphase (20 min) (A) or prolonged metaphase (60 min) (B), with corresponding DIC/mCherry images of time lapse on the left and F-actin staining on the right. Scale bar: 10 μm. Also see Movie S5. (C) Graph plots pMLC mean intensity in mitotic cells of different metaphase duration as indicated. From left to right, n=17, 67, 75, 26, respectively. Mean pMLC intensity of individual cell was calculated by normalization of background-subtracted pMLC intensity over cell area. Data are mean ± SD of 20 fields with totally more than 10 thousands of cells analyzed. (D) The RhoA activity dynamics over metaphase measured by FRET analysis in 21 mitotic cells depleting CDC20. All background-subtracted mean FRET intensities were normalized over their corresponding mean FRET intensities at the time points of nuclear envelop breakdown (NEB) or cell adherence (AD) right before rounding up, whichever applied. Cn refers to different cell analyzed. Also see Movie S6. (E) Quantification of CIC structures in cells overexpressing RhoA. Data are mean ± SD of 4 or more fields with more than 3000 cells analyzed each. p<0.001. (F) Inhibition of CIC formation in CDC20 depleted cells by Y27632, a ROCKs inhibitor that blocks RhoA signaling. Data are mean ± SD of 4 or more fields with more than 5000 cells analyzed each. *p<*0.0001. (G) Polarized distribution of pMLC at the rear cortex of internalized cell in intermediate CIC structure. Scale bars: 10 μm. Also see Figure S4E. (H) Rnd3-GFP localizes at the cell-cell junction during CIC formation. Upper graph shows line scan analysis, channeled images were shown underneath. Scale bar: 10 μm. (I) Rnd3 mRNA level examined by quantitative PCR (qPCR) upon knockdown by RNA interference. Data are mean ± SD of triplicate experiment. *p<*0.001. (J) Representative images for junctional localization of pMLC2 (red) MCF10A cell doublets (right panel) upon Rnd3 depletion (Rnd3 KD), E-cadherin (green) staining indicates cell junctions. Scale bars: 10 µm. Arrowheads indicate pMLC2 staining at cell junctions. Also see Figure S4G (K) Quantification of mintotic CIC formation in control (siNC) and Rnd3 (siRnd3) depleted cells. data are mean ± SD of 4 or more fields with more than 5000 cells analyzed each. *p*<0.01.

### P53-regulated Rnd3 is Essential for Compartmentalizing RhoA Activities and Mintosis

Since polarized distribution of actomyosin, the downstream effector of RhoA signaling, is essential for cells’ penetration into their neighboring cells (Sun *et al.*, 2014a), we examined the expression of p-MLC, the readout of contractile actomyosin and RhoA activity, in the intermediate mintotic CIC structures. As shown in Figure 5G and S4E, p-MLC displays typical asymmetric distribution pattern, characterized by higher intensity at the rear cortex than that at cell-cell junctions marked by E-cadherin where RhoA activities were actively suppressed by inhibitors such as p190A RhoGAP (Sun *et al.*, 2014a). While wild type p53 was known to negatively regulate RhoA signaling through its downstream target Rnd3 (Ongusaha *et al*, 2006; Zhu *et al*, 2014), an atypical Rho GTPase that inhibits RhoA signaling by directly targeting ROCK I (Ongusaha *et al.*, 2006; Riento *et al*, 2003) and p190A RhoGAP (Wennerberg *et al*, 2003). We therefore hypothesized that Rnd3 may facilitate CIC formation via helping compartmentalize RhoA activities. In agreement with this notion, Rnd3, regulated by p53 (Figure S4F), specifically localizes at cell-cell junctions while MLC is enriched at the cortical periphery away from junctions during CIC formation (Figure 5H). SiRNA-mediated Knockdown of Rnd3 led to increased pMLC staining at cell-cell junctions (Figure 5I-J and S4G), suggesting that the junctional-localization of Rnd3 spatially restricts RhoA pathway activity and inhibits myosin contraction at cell-cell junctions, which promotes CIC formation. Consequently, Rnd3 depletion significantly inhibited mintotic CIC formation (Figure 5K). Together, the above data fits a model whereby p53-regulated Rnd3 compartmentalizes RhoA activity in daughter cells from prolonged mitosis to drive CIC formation.

### Mintosis Selectively Targets Mitotic Progenies of Non-Diploidy

Prolonged metaphase due to SAC activation allows cells to fix problems during chromosome segregation. Nevertheless, prolonged mitosis also elicit structural DNA damages (Dalton *et al*, 2007; Ganem & Pellman, 2012) that, via truncated DNA damage response (DDR), induce whole-chromosome missegregation (Bakhoum *et al*, 2014), leading to non-diploid daughter cells (Bakhoum *et al*, 2017). We therefore hypothesized that mintosis was set to ensure selective elimination of those non-diploid cells. Fluorescent in situ hybridization (FISH) was employed to access ploidy changes in MCF10A cells plated on gridded-glass bottom dish following 24 hour-time lapse microscopy, and the metaphase and position histories of target cells were determined by retroactive review of the time lapse records (Movie S7). While mitosis of short metaphase, based on the probes used, generally gave rise to diploid progenies (96.9%) (Figure 6A and 6C), cells of prolonged metaphase penetrating their neighbors to form CIC structures were largely non-diploid (50.7%) (Figure 6B, 6D and S5A). This non-diploid rate of mintotic inner cells is similar to that (43.9%) of inner cells of pre-existing CIC structures (Figure 6E), consistent with our observation that majority of CIC structures were from prolonged mitosis. Intriguingly, the sibling cells that did not participate into CIC formation were largely diploid (88.5%) (Figure 6F) in a rate similar to the whole MCF10A population (89.9%) (Figure 6G). And a small portion of inner cells that eventually got released from CIC structures were also largely diploid (83.8%) (Figure 6H and Figure S5B). Therefore, mintosis selectively targets non-diploid daughter cells for elimination, which may play an essential role in maintaining population genetic integrity, as blocking mintosis by Y27632, an inhibitor of ROCK kinases prerequisite for CIC formation (Sun *et al.*, 2014b), significantly increased non-diploid cells in the cell population (Figure 6I). Thus, our data support that p53-regulated mintosis may serve as a cellular mechanism of mitotic surveillance.

**Figure 6.**
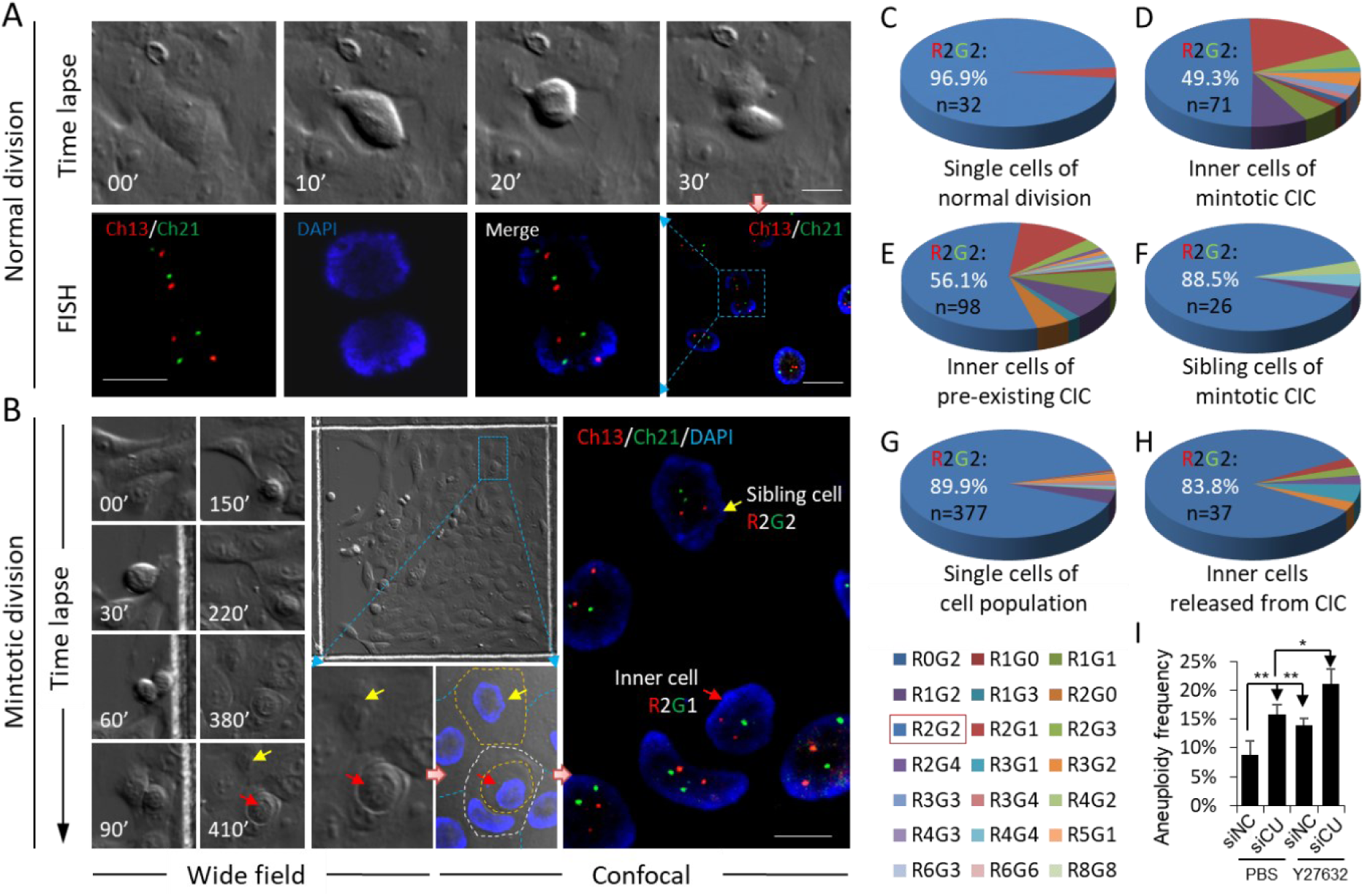
Mintosis selectively targets non-diploid cells for CIC-mediated death. (A) Representative images showing FISH result of normal cell division. Upper panel shows DIC image sequence of time lapse. Lower panel shows FISH results in two daughter cells. Scale bar: 20 μm for right, 10 μm for left. Also see Movie S7. (B) Representative images showing FISH result of cell division leading to CIC formation. Left panel shows DIC image sequence of time lapse. Middle panel shows the positional information of target cells/structures in gridded glass bottom dish at the end of time lapse imaging. Right panel shows FISH results of selected region in middle panel. Scale bar: 10 μm. Yellow arrow indicates adherent sibling daughter cell, red arrow indicates daughter cell internalized to form CIC structure. FISH results are presented as RnGn, R for red probe, G for green probe, n for probe number. Also see Movie S7. (C-H) Detail quantification and classification of FISH results for daughter cells of normal division with metaphase not more than 30 min (C), daughter cells from cell division that were internalized to form CIC structures (D), inner cells of pre-existing CIC structures of unknown origin (E), daughter cells from cell division that did not form CIC structures (F), single cells of unknown origin in the cell population (G), and internalized cells that were finally released from CIC structure (H), see Figure S5B and Movie S7. FISH results are presented as RnGn, R for red probe, G for green probe, n for probe number. The percentage of R2G2 together with cell number analyzed were shown for each pie picture. The related histories for all cells analyzed were determined by time lapse imaging. (I) Quantification of non-diploid cells following mintosis blockade. siNC: non-target control siRNA; siCU: siRNA for CUEDC2. “*” for p<0.05; “**” for p<0.01. For each condition, quantification was performed on >400 cells..

## Discussion

In summary, we proposed a CIC-mediated mechanism of post-mitotic surveillance as mintosis whereby non-diploid progenies from prolonged mitosis were eliminated in a p53-dependent manner. In this model (Figure 7), γH2AX-marked DNA damages associated with prolonged mitosis in mother cells activate p53 pathway in daughter cells that are non-diploid due to chromosome missegregation, p53 upregulates the expression of its downstream target Rnd3 which locally inhibits RhoA signaling and actomyosin contraction at cell-cell junctions, leading to asymmetric RhoA activation at the rear cortex of daughter cells to drive cell internalization and subsequent non-apoptotic death. Thus, this work uncovered a novel non-apoptotic mechanism by mintosis for p53 to maintain genome integrity.

**Figure 7.**
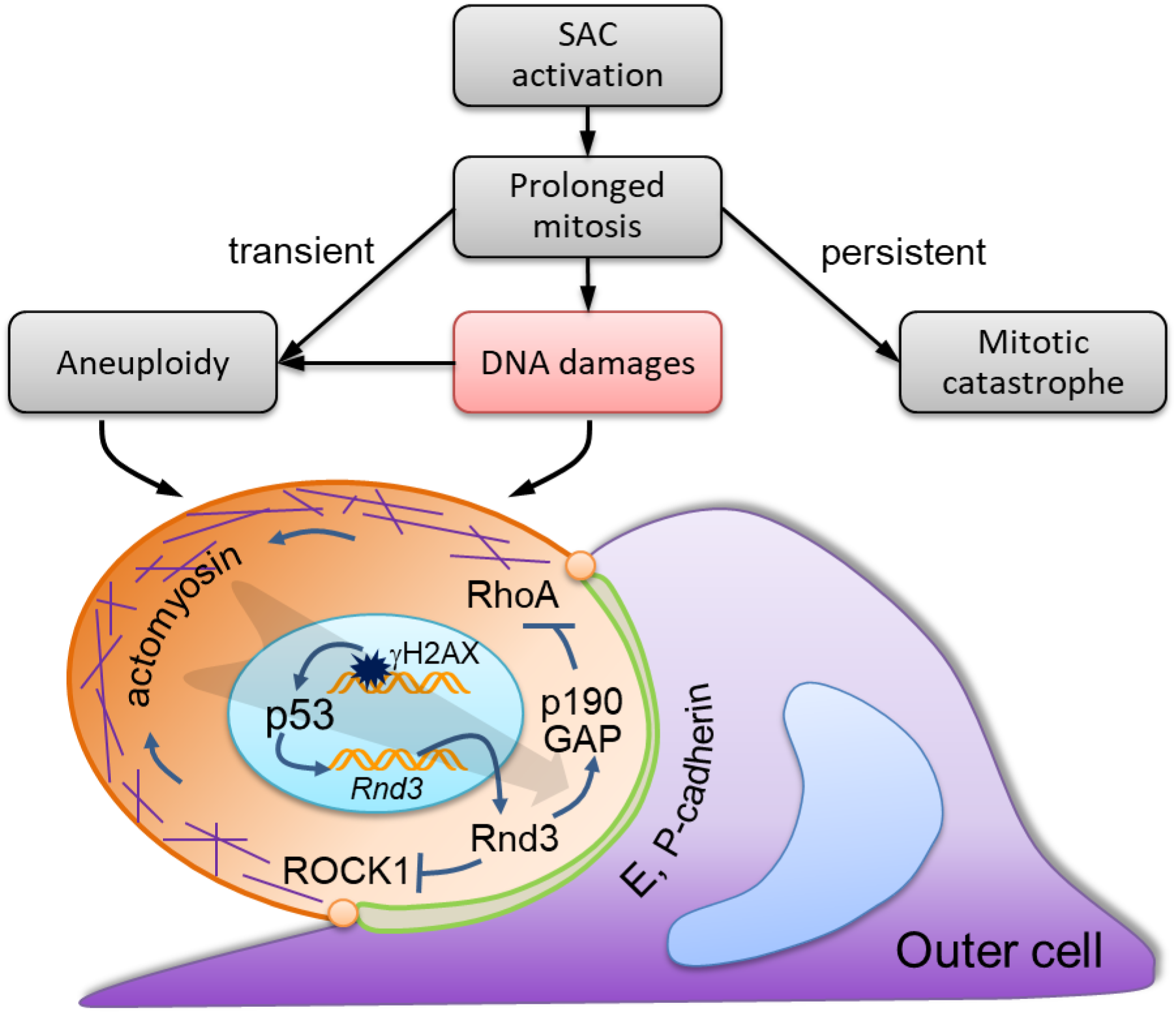
Working model for p53-depedent postmitotic surveillance by mintosis. SAC activation leads to mitotic arrest characterized by prolonged metaphase. While persistent mitotic arrest activates cell death by mitotic catastrophe prior to cytokinesis, transient mitotic arrest may cause DNA damages that promote chromosome missegregation to give rise to daughter cells of aneuploidy. DNA damages marked by gH2AX activate DDR signaling and p53 in daughter cells. p53 upregulates the expression of downstream target Rnd3 which, by targeting ROCK1 and p190 RhoGAP, inhibits RhoA activities at cell-cell junction where lie E-, P-cadherin, leading to polarized activation of RhoA-actomyosin at rear cortex which drives cell internalization to form CIC structures. Subsequent death promoted by the outer cell eliminates those aneuploid daughter cells to maintain genome integrity.

DNA damages associated with mitosis are generally phenotyped as prolonged mitosis or mitotic arrest, which could also cause DNA damages (Dalton *et al.*, 2007; Ganem & Pellman, 2012). Formation of this vicious loop is largely due to impaired DNA damage response (DDR) which, though being capable of sensing DNA damages, likely stops short of downstream damage repair pathways (Heijink *et al*, 2013). Though activation of this partial DDR was believed to prevent fusion of exposed telomeres under normal circumstance (Orthwein *et al*, 2014), it left unrepaired DNA damages, pre-mitotic or acquired, into mitotic progenies under stressed conditions (Denchi & Li, 2014). Moreover, activation of the partial DDR during mitosis, instead helps fix errors in DNA, actually could result in severe chromosome missegregation, which eventually gives rise to aneuploid progenies (Bakhoum *et al.*, 2017; Bakhoum *et al.*, 2014). Therefore, surveillance mechanisms are critical to deal with daughter cells from prolonged mitosis. Previous studies indicated that prolonged mitotic progenies might be surveilled by either growth arrest or apoptosis (Joerger & Fersht, 2016; Lambrus & Holland, 2017; Vitale *et al.*, 2011). Whereas in this study, we identified a novel surveillance mechanism executed by CIC-mediated non-apoptotic death termed mintosis. Although aslo being activated by prolonged mitosis via p53-dependent pathway, mintosis likely represents a unique process independent from growth arrest and apoptosis because depleting p21, a p53 downstream effector that is required for growth arrest/senescence, didn’t block mintotic CIC formation at all (data not shown), and depletion of Rnd3, a p53 target gene that was identified as suppressor of ROCK I-mediated apoptosis (Ongusaha *et al.*, 2006; Paysan *et al*, 2016), significantly suppress mintosis (Figure 5I-K).

Interestingly, while our work demonstrates a positive role of CIC formation in maintaining genome stability by eliminating aneuploidy cells, previous work supported entosis as a cellular mechanism to promote genome instability by inducing aneuploidy (Krajcovic *et al.*, 2011). This clear conceptual discrepancy is actually due to different cellular and molecular contexts where CIC formation may work. At cellular level, Krajcovic *et al* found that the presence of inner cell could physically block cytokinesis of outer cell, leading to outer cell binucleation and subsequent aneuploidy of its offspring following next round of cell division (Krajcovic *et al.*, 2011). This effect may be propagated to promote genome instability in the context of tumor cells that are tolerated to aneuploidy. This idea was supported by recent work in lung cancer cells Mackay *et al* (Mackay *et* al., 2018). Whereas this work investigates CIC’s role in a non-tumor context, in which aneuploid daughters from prolonged mitosis were internalized as inner cells and eliminated. Thus, it’s likely that CIC formation affects genome stability differentially via inner cell and outer cell, respectively, depending on cells’ tolerance to aneuploidy. For tumor cells who are generally aneuploidy-tolerant, CIC formation may promote genome instability by inducing cytokinesis failure of outer cells; for non-transformed epithelial cells, CIC formation may function to counteract genome instability by internalizing and eliminating aneuploid cells. Of note, since aneuploidy is toxic and lethal to non-transformed cells, the aneuploid progenies of binucleated outer cells wouldn’t survive for long period (data not shown). This is also true for tumor cells that are not tolerant to aneuploidy (Mackay *et al.*, 2018). Therefore, CIC formation is genome instability- or tumor-suppressive anyway when cells are not tolerant to aneuploidy. At molecular level, CIC’s effects on aneuploidy and genome stability are controlled by p53 status. Mackay *et al* found that only mutant p53, but not wild type or null p53, could promote genome instability via CIC formation by conferring cancer cells winner/outer identity and allowing survival and population of their aneuploidy progenies (Mackay *et al.*, 2018). Whereas, our work demonstrated that wild type p53 endowed loser/inner identity to cells during CIC formation by facilitating the establishment of polarized RhoA activities. Together, we propose that CIC formation, depending on p53 status, may play dual roles in genome instability and tumorigenesis as well. For cancer cells with mutant p53, CIC formation plays promotive role by facilitating selection of cancer cells harboring mutant p53; for normal epithelial cells or cancer cells that contain wild type p53, CIC formation play suppressive role by eliminating aneuploidy cells expressing wild type p53 through mintosis. It’ll be of interest to explore whether other oncogenic mutations may also regulate CIC’s functional outcomes.

Recently, two other works reported entosis induction in confined contexts in adherent monolayer cultures (Durgan *et al.*, 2017; Hamann *et al*, 2017). One of work demonstrated that glucose starvation, an extreme biological condition, was capable of promoting CIC formation mainly in the context of cancer cells like MCF7 in an AMP activated protein kinase (AMPK)-dependent way (Hamann *et al.*, 2017). Interestingly, although AMPK is a well-known energy sensor, it was found in mitotic apparatus and involved in the regulation of mitotic progression and completion (Banko *et al*, 2011; Li & Zhang, 2017). So we hypothesize that glucose induced activation of AMPK may influence normal mitosis, which triggers mintosis and contributes to CIC formation in adherent cells that is yet to be validated. Another work reported that depletion of CDC42, a polarity protein, induced CIC formation in 16HBE cells (Durgan *et al.*, 2017). Although mitosis was found linked to CIC formation, the work failed to identify the essential role of mitotic arrest and DNA damages in CIC formation, hence, couldn’t tell why most mitosis failed to give rise to CIC structures and why reagents blocking mitosis unexpectedly induced CIC formation. Intriguingly, similar to work by Wan et al (Wan *et al*, 2012), where depletion of polarity protein PAR3 resulted in enhanced formation of CIC structures in adherent MDCK cells, polarity changes were not regarded as the reason for enhanced CIC formation (Durgan *et al.*, 2017; Wan *et al.*, 2012). Whereas, we found that depleting polarity protein such as CDC42 in MCF10A and tumor cells could promote CIC formation that was also preceded by prolonged mitosis (data not shown), we therefore speculate that altered mitosis might be a shared route for multiple factors, such as glucose starvation and depleting polarity proteins and the like, to activate mintosis that works in multiple contexts to maintain homeostasis.

## Materials and Methods

### Cell culture and constructs

MCF7 and 293FT cells were maintained in Dulbecco’s modified Eagle’s medium (DMEM) supplemented with 10% fetal bovine serum (PAN-Biotech). MCF10A and its derivatives were cultured in DMEM/F12 supplemented with 5% horse serum (GIBCO, #16050-122), 20 ng/ml EGF (Peprotech, #96-AF-100-15-100), 10 µg/ml insulin (Sigma, I-5500), 0.5 µg/ml hydrocortisone (Macgene, CC103), and 100 ng/ml cholera toxin (Sigma, C8052). pBabe-H2B-mCherry was a gift from Dr. Michael Overholtzer. pBabe-RhoA biosensor was a gift of Dr. Klaus Hahn from Addgene (12602). pLKO-shp53 was a gift of Dr. Bob Weinberg from Addgene (19119). GFP-p53 was a gift of Dr. Tyler Jacks from Addgene (12091). Expression plasmid for GFP-Rnd3 was constructed by inserting synthesized human Rnd3 ORF into pQCXIP-GFP vector by *Xho* I and *Bam*H I sties. Expression plasmid for MLC-mCherry was constructed by inserting synthesized chicken MLC ORF into pBabe-mCherry vector by *Xho* I and *Bam*H I sties.

### Antibodies and chemical reagents

Antibodies with working dilution factors, company source and catalog number include: anti-pMLC (1:200; Cell Signaling; #3671), anti-E-cadherin (1:200; BD Biosciences; 610181), anti-γH2AX (1:200; Cell Signaling; #9718 for IF; 1:500; ABclonal; AP0099 for WB), anti-H2AX (1:1000; ABclonal; AP0823), anti-pATM (1:500; Boster; BM4008), anti-ATM (1:1000; Proteintech; 27156-1-AP), anti-pp53 (1:500; Cell Signaling; #9386), anti-p53 (1:1000; Santa Cruz; sc-126), anti-Rnd3 (1:1000; Sino Biological; 101056-T32). Secondary antibodies include Alexa Fluor 568 anti-mouse (1:500; Invitrogen; A11031), Alexa Fluor 568 anti-rabbit (1:500; Invitrogen; A11036), Alexa Fluor 488 anti-mouse (1:500; Invitrogen; A11029) and Alexa Fluor 488 anti-rabbit (1:500; Invitrogen; A11034). Alexa Fluor®647 Phalloidin (1:200; Invitrogen; A22287). DAPI was purchased from Sigma (D8417). ROCK inhibitor Y27632 was purchased from TOCRIS (1254) and used at final concentration of 10 µM. Ro-3306 (HY-12529), a potent and selective inhibitor of CDK1, and SU-9516 (HY-18629), a selective CDK2 inhibitor, were purchased from MedChem Express. DNA damage inducers Mitomycin (T6890) and Bleomycin (T6116), and inhibitors for ATM (T2474), ATR (T3338), CHK1 (T2033), CHK2 (T7080) were purchased from TargetMol. Collagen Type I is a product of BD Biosciences (#354236).

### Virus production and infection

Stable expression cell lines were established by virus infection. Briefly, 1×10^6^ 293FT cells were plated into 6-well plate coated with collagen I (BD Bioscience, #354236), transfection was performed with retroviral constructs together with packaging plasmids, and viruses were collected twice at 24 h intervals. To infect cells, cells were cultured in 1 ml viral supernatant mixed with 1 µl polybrene of 10 µg/ml stock for 6 h followed with regular media. Cells were selected with appropriate antibiotics (2 µg/ml puromycin or 400 µg/ml G418 for MCF10A, 1µg/ml puromycin for MCF7) for 7 days.

### RNA interference

siRNAs were from GenPharma (Shanghai, China). For individual siRNA trans fection, cells (1×10^5^/well) were plated into 12-well glass bottom plate and cultured overnight, then transfected with 50 nM siRNA using Lipofectamine® RNAiMAX (Invitrogen, #13778-150). Cell s were fed with fresh full media 6 h later. siRNA sequences: CUEDC2: 5’-CAUCAGAGGAGAA CUUCGA-3’; CDC20: 5’-CCACCAUGAUGUUCGGGUATT-3’; MAD2 : 5’-GGAAGAGUCGG GACCACAGTT-3’; BuBR1: 5’-CGGGCAUUUGAAUAUGAAATT-3’; ESPL1: 5’-GCUUGUGA UGCCAUCCUGATT-3’; H2AX-1: 5’- GGGACGAAGCACUUGGUAACA-3’; p53-1: 5’- AAGA CUCCAGUGGUAAUCUAC-3’; Rnd3-1: 5’- GAUCCUAAUCAGAACGUGAAA-3’; Rnd3-2: 5’- AUCCUAAUCAGAACGUGAAAU-3’; ATM: 5’- GCCUCCAAUUCUUCACAGUAA-3’; ATR: 5’- GAUGAACACAUGGGAUAUUUA-3’; CHK1-1: 5’- GUGACAGCUGUCAGGAGUAUU-3’; CHK1-2: 5’- GCCCACAUGUCCUGAUCAUAU-3’; CHK2-1: 5’- GAACAGAUAAAUACCGAA CAU-3’; CHK2-2: 5’- CGCCGUCCUUUGAAUAACAAU-3’; Negative Control: 5’-UCUCCGAA CGUGUCACGUTT-3’.

### Reverse transcription-quantitative PCR (RT-qPCR)

Total RNA was isolated from cells 48 h after siRNA transfection using TRIzol reagent (Invitrogen, #15596026). One microgram of total RN A was converted into cDNA using *TransScript*^®^ One-Step gDNA Removal and cDNA Synthesis S uperMix (Transgen Biotech, #AT311-02) according to manufacturer’s instruction. The quantitative PCR (qPCR) was performed on 15 ng of cDNA from each sample using SYBR Green Real-time PCR Master Mix (TOYOBO, #QPK-201) based on the recommendations of manufacturer. Primers pairs spanning at least two exons were confirmed by NCBI Primer-BLAST: CUEDC2: 5’-TGAG CGATGCCAGGAACAA-3’ and 5’-CTCCTCCTCAGCGCCAGTT-3’; CDC20: 5’-TTCCCTGCC AGACCGTATCC-3’ and 5’-CAGCCAAGTAGTTGCCCTC-3’; MAD2: 5’-TTCTCATTCGGCAT CAACA-3’ and 5’-TCTTTCCAGGACCTCACCA-3’; BuBR1: 5’-TCTTCAGCAGCAGAAACGG -3’ and 5’-TCATTGCATAAACGCCCTA-3’; ESPL1: 5’-CCCCACTTCGGGCATTGTA-3’ and 5’- GGGCAAAGTCATAAACCACC-3’; BUB1: 5’-AGAAATACCACAATGACCCAA-3’ and 5’-AG GCGTGTCTGAAATAACC-3’; H2AX: 5’- CCCTTCCAGCAAACTCAACTCG-3’ and 5’- AAAC TCCCCAATGCCTAAGGT-3’; p53: 5’- ACCACCATCCACTACAACTACAT-3’ and 5’- CTCCC AGGACAGGCACAAA-3’; Rnd3: 5’- TCTTACCCTGATTCGGATGC-3’ and 5’- TCTGACGCTA TTTTCCGACT-3’; ATM: 5’- GCACAGAAGTGCCTCCAATTC-3’ and 5’- ACATTCTGGCACG CTTTG-3’; ATR: 5’- GCCGTTCTCCAGGAATACAG-3’ and 5’- GAGCAACCGAGCTTGAGAG T-3’; CHK1: 5’- GGATGCGGACAAATCTTACCA-3’ and 5’- CCTTAGAAAGTCGGAAGTCAA CC-3’; CHK2: 5’- GTCATCTCAAGAAGAGGACT-3’ and 5’- GAGCTGTGGATTCATTTTCC-3 ‘; HPRT: 5’-AGGCCATCACATTGTAGCCCTCTGT-3’ and 5’-TACTGCCTGACCAAGGAAAG CAAAGT-3’. The PCR reactions run on the following conditions: initial denaturing at 95°C for 30 sec, followed by 35–40 cycles of 95°C for 5 sec, 60°C for 10 sec and 72°C for 15 sec, melting cur ves were examined at 37°C for 30 sec before cooling. Each result was from three independent biological replicates for all analyses performed in this work. The qPCR results were analyzed using 2^-ΔΔCT^ method and presented as relative quantity of transcripts with HPRT as the reference gene.

### Cytospin and entotic CIC quantification

Described protocol (Sun & Overholtzer, 2013) was slightly modified to examine cells abilities to form CIC structures. Briefly, cells were cultured in suspension for 6 h in 6-well plate pre-coated with 1 ml solidified soft agar (0.5%) and then mounted onto glass slides for 3-min centrifugation at 400 rpm to make cytospin. Cells were fixed by 4% PFA and immunostained with E-cadherin antibodies followed by mounting with Antifade reagent with DAPI. CIC structures with more than half of cell body internalized were counted.

### Time lapse imaging and mintotic CIC quantification

Wide field imaging was performed on cells plated in glass bottom dish or plate (Nest Biotechnology Co.) by Nikon Ti-E microscope equipped with motorized stage and Neo Vacuum cooled Scientific CMOS Camera (Andor Technology). Images were collected every 10 or 15 min for 24 h using 10x or 20x Apo objective lens with 15 ms exposure for DIC channel and 150 ms exposure for mCherry channel. Cells were cultured in humidified chamber supplied with 5% CO_2_ at 37°C during imaging. Image sequences were reviewed using Nikon NIS-Elements AR 4.5 software. Mitotic entry was judged morphologically by either condensed chromatin as indicated by H2B-mCherry condensation or cells’ round up. Mitotic anaphase was judged by chromosome separation labeled with H2B-mCherry. The duration of metaphase was determined from the first frame of mitotic entry to the first frame of mitotic anaphase. CIC structures were determined morphologically by complete enwrapping of cells into their neighbors, typically with a crescent nucleus in the outer cells. Total cells in each field were counted on mCherry-postive nuclei. CIC frequency was presented as CIC number divided by total cells in each field.

For CIC formation induced by DNA damages, MCF10A/H2B-mCherry cells in 12 well plate (2.5*10^4/well) were synchronized by 100ng/m nocoldazole for 6 h followed by treatment of DNA damage inducers mitomycin (0.5 μM, 1 μM, 2 μM) for 3.5 h. Then, drugs were washed out for 20 h time lapse in full medium. CIC formation was quantified as did above.

### Time lapse-associated FISH (fluorescence in situ hybridization)

Prior to FISH, MCF10A cells of 3×10^5^ were first cultured in gridded glass bottom dish (µ-Dish 35mm Grid-500, ibidi; #81168) for 20 h, followed by time lapse microscopy with images collected every 10 min for 16 h by using 10x Apo objective lens in DIC channel. Then, cells were immediately fixed at room temperature for 20 min with freshly prepared solution (methanol/glacial acetic acid = 3:1) after briefly washed twice with PBS. Fixed cells were initially baked at 56 °C for 60 min, subsequently incubated in 100 µg/ml RNase (pH=7.0±0.2) for 1 h and then 20 mg/ml pepsin-0.01 M HCl for 10 min at 37 °C. After washed twice with 2× SSC at room temperature for 5 min, samples were dehydrated in 70%, 85%, and 100% pre-cooled ethanol for 2 min respectively, and then air-dried. Hybridization was performed with two-probe FISH kit (F01010-00, GP Medical Technologies, Ltd) following the manual provided. Briefly, sample was denatured at 75°C for 10 min and then preceded to hybridization at 42°C for 16 h in a humidified cassette. Following hybridization, sample was serially washed with 0.4x SSC containing 0.3% NP-40 (pH=7.0±0.2) at 65 °C for 3 min, 2x SSC containing 0.1% NP-40 (pH=7.0±0.2) at room temperature for 1 min and finally 70% ethanol at room temperature for 3 min before counterstained with DAPI in darkness for 10-15 min and mounted. Images were taken by using *Ultraview Vox* spinning disc confocal system (Perkin Elmer) equipped with a Yokogawa CSU-X1 spinning disc head and EMCCD camera (Hamamatsu C9100-13) on Nikon Ti-E microscope. Analysis was performed with Volocity software (Perkin Elmer). Information on cell position, division and CIC formation were determined based on time lapse imaging and grids on the glass bottom.

### Time lapse-associated immunostaining (TLAS)

For phospho-Myosin Light Chain 2 (pMLC) staining in mitotic cells, MCF10A/H2B-mcherry cells in gridded glass bottom dish (µ-Dish 35mm Grid-500) were transfected with CDC20 siRNA as described above. Time lapse imaging was performed by 10x Apo objective lens in DIC and mCherry channels next day, with images captured every 10 min for 4 h before fixing with 4% PFA and preceded to routine staining. Briefly, the fixed sample was permeabilized with 0.2% Triton-X 100/PBS for 5 min and blocked with 5% BSA for 1 h before incubated with primary antibody at 4°C overnight followed fluorophore-labeled secondary antibody, cells were then co-stained with phalloidin and DAPI for 20 min and then mounted with Prolong Gold antifade reagent (Invitrogen). Images were captured with Nikon Ti-E microscope equipped with Neo Vacuum cooled Scientific CMOS Camera (Andor Technology) and analyzed with Nikon NIS-Elements AR 4.5 software. Mean pMLC intensity of individual cell was calculated by equation: (pMLC-background)/area. TLAS of γH2AX and p53 were performed following the protocol for pMLC with slight modification that time lapse was performed for 20 h in control or CUEDC2-depleted MCF10A cells.

### Immunostaining and Immunoblotting

Cultured cells were stained following protocol above in TLAS. For tissue sections, samples were first deparaffinized and antigen retrieved following routine procedures and then preceded to immunostaining as did in fixed cells above. Confocal images were captured and processed by *Ultraview Vox* confocal system (Perkin Elmer) on Nikon Ti-E microscope. Immunoblotting was performed following standard precedures, briefly, protein samples were separated by SDS-PAGE and then transferred onto PVDF membrane, where specific antibodies were used to probe target proteins.

### FRET

Briefly, activation levels of RhoA were measured by monitoring the ratio of ECFP to Citrine-YFP FRET and ECFP intensities (Pertz *et al*, 2006; Sun *et al.*, 2014a). Images were acquired on a Nikon Ti-E inverted microscope using a Neo Vacuum cooled Scientific CMOS Camera (Andor Technology) mounted on the bottom port with a set of excitation/emission filter wheels to direct the DIC, ECFP, FRET, and Citrine-YFP signals sequentially. Images were obtained using a Nikon 20x/0.75 CFI Plan Apochromat Lambda lens and Nikon NIS-Elements AR 4.5 software. The filter sets used for ratiometric imaging were (Excitation, emission, respectively, Chroma Technology): ECFP: ET438/24, ET482/35; FRET: ET438/24, ET540/30; and Citrine-YFP: ET513/17, ET540/30. Cells were illuminated by LUMENCOR SPECTRA X Light Engine with 438/24 and 513/17 excitation filters. CFP, FRET and DIC images were recorded with 1 × 1 binning. The FRET module of Nikon NIS-Elements AR 4.5 software was used to process image sequences. The background-subtracted images from two cameras were aligned to ascertain optimal registration with subpixel accuracy. A linear rainbow pseudocolor lookup table was applied to the ratiometric images.

### Patient samples

Breast cancer sections were obtained from 307 Hospital under the hospital’s regulations and ethics. Sections stained with E-cadherin or HER2 were scanned by NanoZoomer-SQ (Hamamatsu) digital slide scanning system. CIC structures were judged by fully enclosing of one or more cells within another cell based on membrane contour line. Patient sections were first screened based on the number of CIC structures in fields of 40x magnification, those that had less than 1 CIC structures in more than 3 fields were scored as low CIC, more than 15 CIC structures in 3 fields as high CIC.

### Statistics

All assays were carried out in triplicate or more. Data were expressed as means with standard deviations (SD) or standard error of mean (SEM). *P*-values were calculated using two-tailed Student’s t-test from Excel or GraphPad Prism software, and *P*-values less than 0.05 were considered statistically significant. Logistic regression analysis in Excel was used to evaluate association between factors.

## Supporting information

Movie S1-CIC formation-s

Movie S2-Mintosis

Movie S3-Inner cell fate

Movie S4-mitotic catastrophe

Movie S5-Division of different metaphase

Movie S6-FRET-one ENTOSIS

Movie S7-FISH

## Acknowledgments

We thank Dr. Overholtzer from Memorial Sloan-Kettering Cancer Center for providing cell lines and reagents. We thank Dr. Louis Hodgson from Albert Einstein College of Medicine of Yeshiva University for assistance in FRET. We thank Dr. Dangsheng Li for discussions and critical reading of the manuscript.

## Funding

This work was supported by the National Key Research & Development Program of China (2016YFC1303303, 2019YFA09003801, 2018YFA0900804), the National Natural Science Foundation of China (31970685, 31671432, 81872314, 31770975, 81572799).

## Author contributions

Q.S. conceived the project and conception. Q.S., H.H. and J.L. designed experiments with the advice from X.N.W. and Z.C.. J.L. performed majority of the experiments with the assistance of X.Y. for RNA interference, phenotype and qPCR, and Z.N. for pathology, immunostaining and FISH. M.W. B.Z. and H.Q helped in protein expression and detection. S.G. and L.G. assisted imaging. Y.T. provided patient samples and related information. Q.S., H.H. and J.L. analyzed the data with the assistance of Y.Z.. Q.S. and J.L. wrote the paper with input from X.N.W, L.M. and H.H., and all authors reviewed the manuscript.

## Competing interests

no conflict of interest to be stated.

## Supplementary Materials

**Figure S1.**
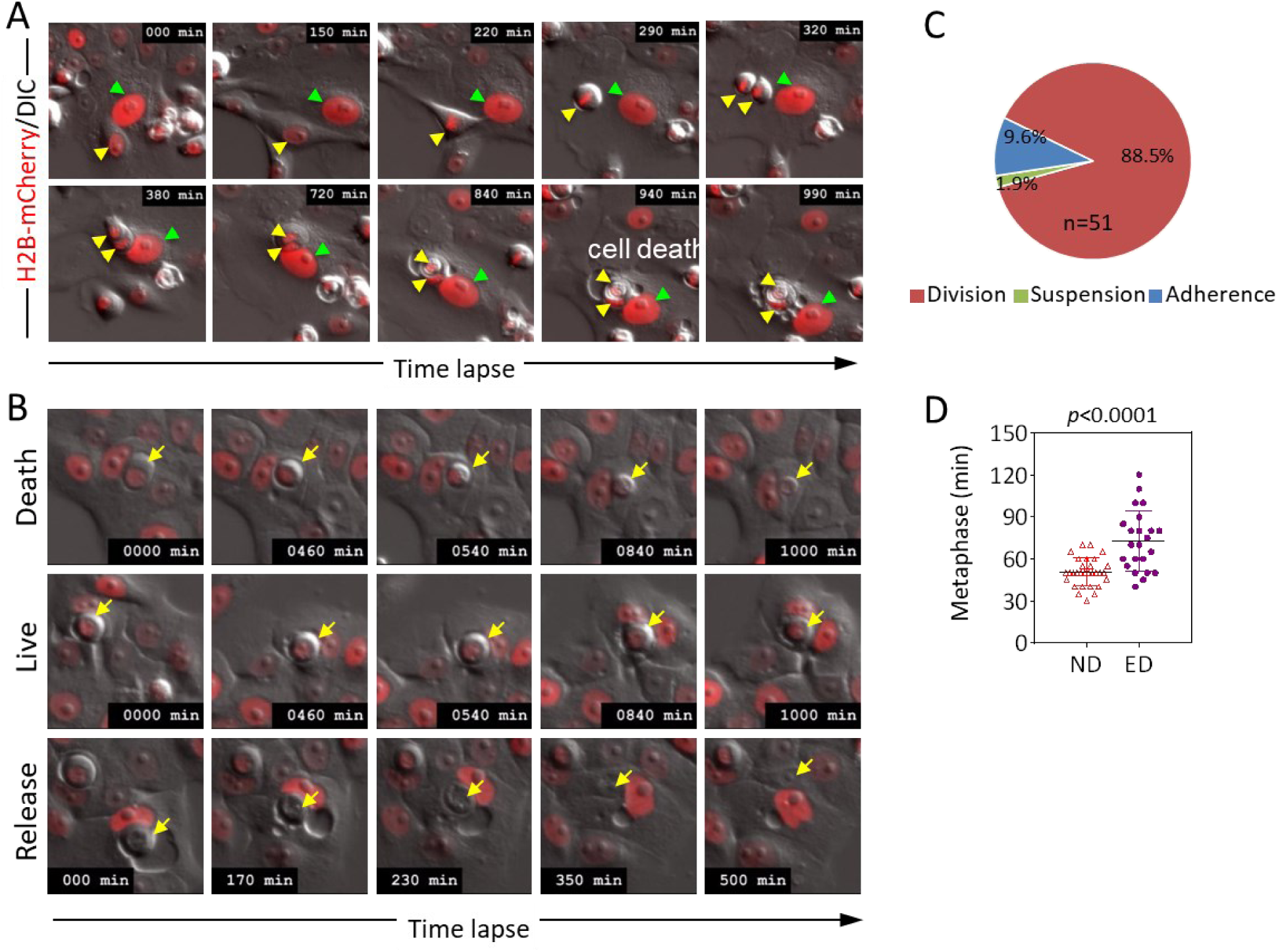
Entosis is coupled with mitotic cell division. Related to Figure 1. (A) Image sequence shows the process of entotic cell-in-cell formation and inner cell death following mitotic cell division in MCF10A/H2B-mCherry cells. Yellow arrows indicate cells undergoing mitosis and internalization, green arrows indicate outer cells. (B) Representative image sequences of different inner cell fates for entotic cell-in-cell structures of MCF10A cells. Arrows indicate inner cells. Also see Movie S2. (C) Quantification of three ways leading to CIC structures in MCF7 cells. n=51. (D) Metaphase analysis of normal cell division (ND) and entotic cell division (ED) referring to cell division leading to entotic CIC formation in MCF7 cells. n=30 for ND, 23 for ED. *p*<0.0001.

**Figure S2.**
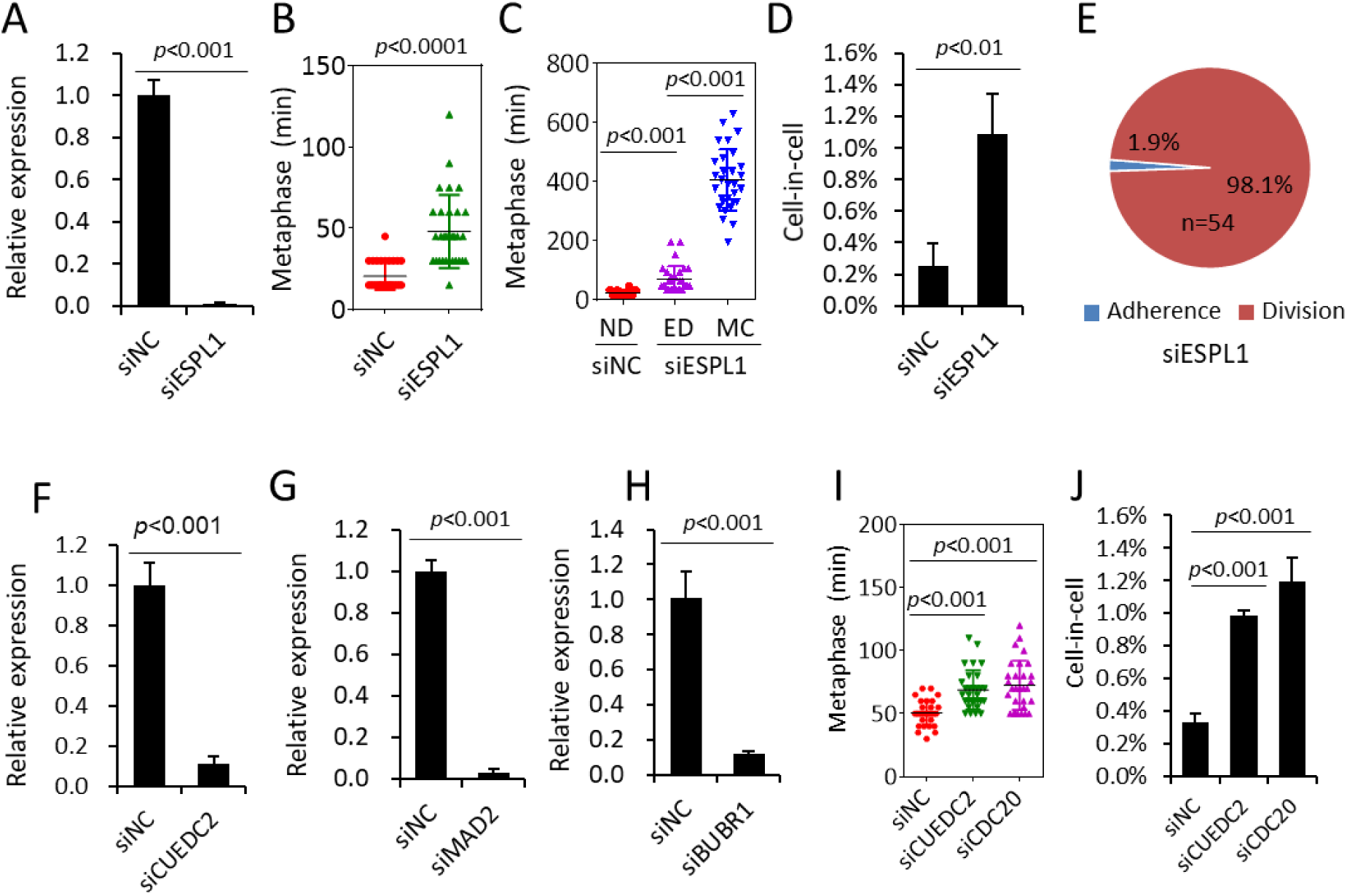
Prolonged mitosis primes cells to undergo entosis. Related to Figure 2. (A) ESPL1 mRNA level examined by quantitative PCR (qPCR) upon knockdown by RNA interference. Data are mean ± SD of triplicate experiment. *p<*0.001. (B) Quantification of CIC structures in control (siNC) and ESPL1 (siESPL1) depleted cells. Data are mean ± SD of 4 or more fields with more than 5000 cells analyzed each. *p<*0.01. (C) Graph plots metaphase duration of control (siNC, n=30) and ESPL1 depleted (siESPL1, n=30) cells that didn’t undergo entosis or mitotic catastrophe. *p<*0.0001. (D) Quantification of CIC formation in control (siNC) and ESPL1 depleted (siESPL1) cells. Data are mean ± SD of 4 or more fields with more than 5000 cells analyzed each. *p<*0.01. (E) Quantification of Entotic Adherence (Adherence) and Entotic Division (Division) in MCF10A cells with ESPL1 depletion. (F) CUEDC2 mRNA level examined by quantitative PCR (qPCR) upon knockdown by RNA interference. Data are mean ± SD of triplicate experiment. *p<*0.001. (G) MAD2 mRNA level examined by quantitative PCR (qPCR) upon knockdown by RNA interference. Data are mean ± SD of triplicate experiment. *p<*0.001. (H) BUBR1 mRNA level examined by quantitative PCR (qPCR) upon knockdown by RNA interference. Data are mean ± SD of triplicate experiment. *p<*0.001. (I) Graph plots metaphase duration of control (siNC) and CDC20 and CUEDC2 depleted (siCDC20, siCUEDC2) MCF7 cells. n=30 for each. *p<*0.001. (J) Quantification of CIC structures in control (siNC) and CDC20 and CUEDC2 depleted (siCDC20, siCUEDC2) MCF7 cells. Data are mean ± SD of 4 or more fields with more than 5000 cells analyzed each. *p<*0.001.

**Figure S3.**
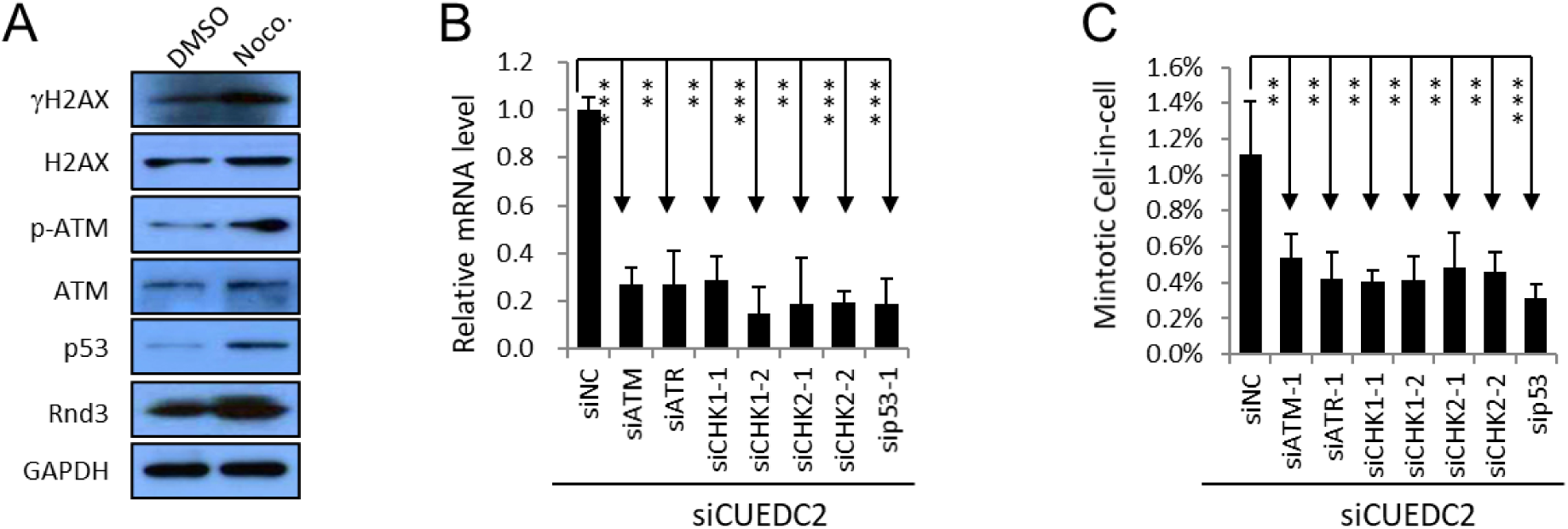
Involvement of DNA damage pathway in mintosis. Related to Figure 3. (A) Activation of DNA damage response (DDR) signaling during mitotic arrest. Expression of genes of DDR pathway in nocodazole (Noco.)-treated MCF10A cells were detected by Western blot. (B) mRNA levels of genes in DDR pathway examined by quantitative PCR (qPCR) upon knockdown by RNA interference. Data are mean ± SD of triplicate experiment. *p<*0.001. (C) Decreased mintotic CIC formation in MCF10A cells upon depletion of genes in DDR pathway by RNA interference. Data are mean ± SD of 4 or more fields with more than 5000 cells analyzed each. “**” for *p*<0.01; “***” for *p*<0.001.

**Figure S4.**
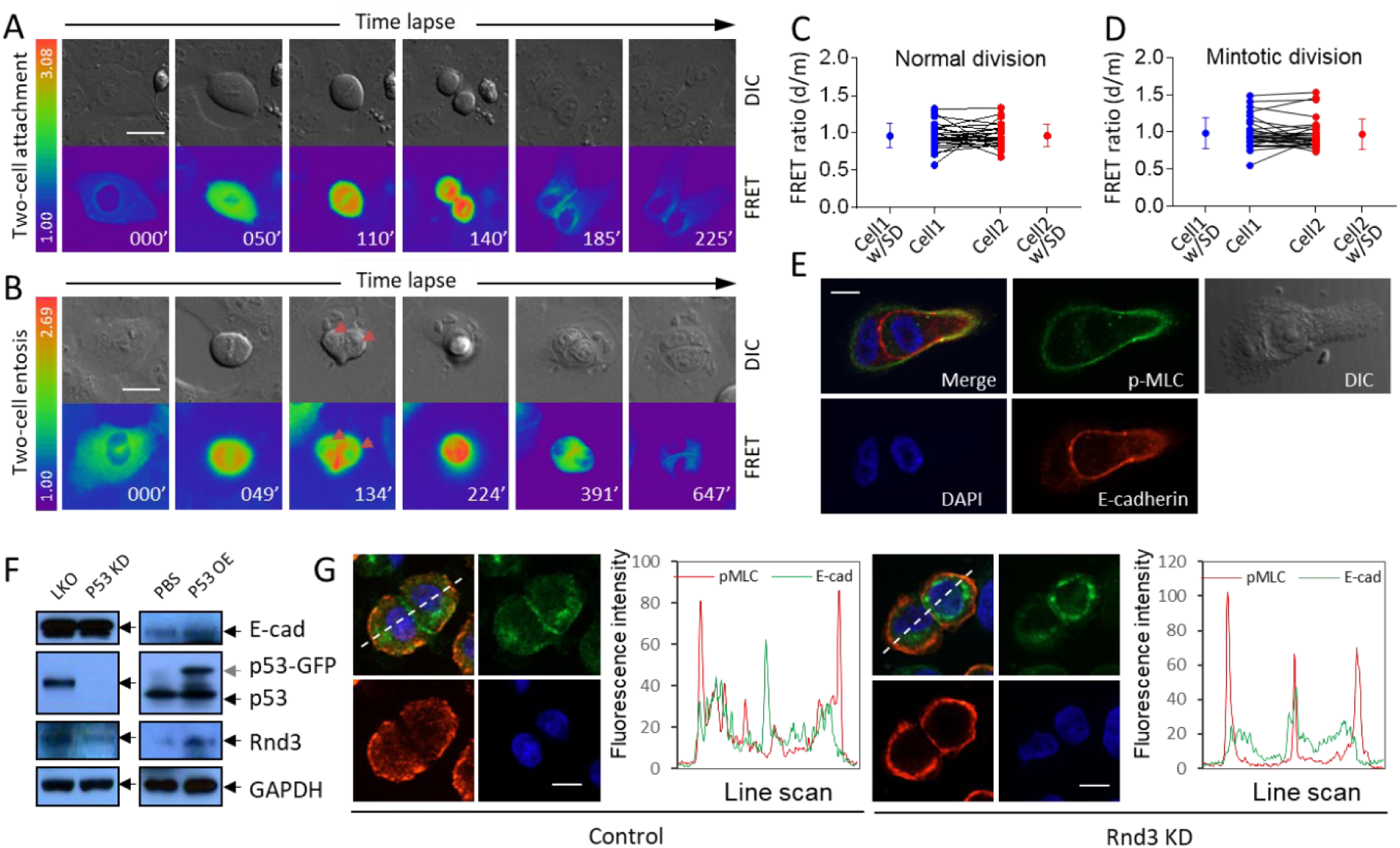
Compartmentalized RhoA activity by Rnd3 is required for mintosis. Related to Figure 5. (A) Representative FRET image sequence shows RhoA activity changes during cell division with two daughter cells attached to plate bottom. Scale bar: 20 μm. (B) Representative FRET image sequence shows RhoA activity changes in cell division with two daughter cells internalized to form CIC structures. Scale bar: 20 μm. (C-D) RhoA FRET ratios between daughter cells right after cytokinesis (d) and their respective mother cells in the end of metaphase (m) in normal cell division (C, n=28) and mintotic cell division (D, n=29). (E) Representative images for polarized distribution of pMLC at the rear cortex of internalized cell in intermediate CIC structure. Scale bars: 10 μm. Identical to Figure 5G. (F) Regulation of Rnd3 expression by p53 in MCF10A cells. Left panel shows decreased Rnd3 expression upon stable p53 knockdown (p53 KD) as compared with control (LKO). Right panel shows increased Rnd3 expression upon p53-GFP overexpression (p53OE). (G) Representative images for junctional localization of pMLC (red) MCF10A cell doublets (right panel) upon Rnd3 depletion (Rnd3 KD), E-cadherin (green) staining indicates cell junctions. Scale bars: 10 µm. Arrowheads indicate pMLC staining at cell junctions. Related to Figure 5J.

**Figure S5.**
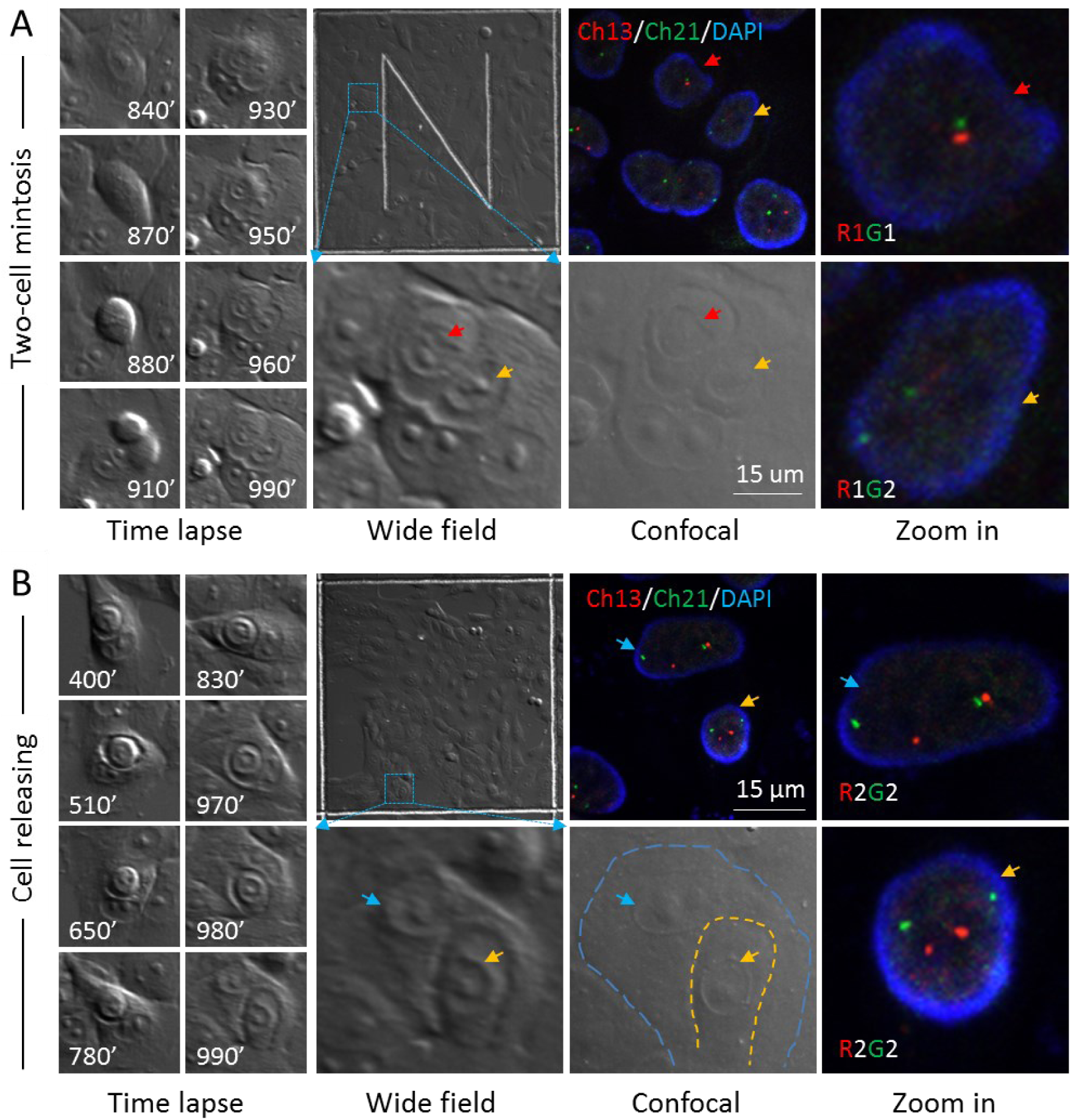
Mintosis selectively targets non-diploid cells for elimination. Related to Figure 6. (A) Representative images showing FISH result of cell division leading to internalization of two daughter cells to form CIC structures. Left panel shows DIC image sequence of time lapse. Middle panel (wide field) shows the positional information of target cells/structures in gridded glass bottom dish at the end of time lapse imaging. Right panels (confocal and zoom in) show FISH results of selected region in middle panel. Scale bar: 15 μm. Yellow and red arrows indicate two daughter cell internalized. FISH results are presented as RnGn, R for red probe, G for green probe, n for probe number. See Movie S7. (B) Representative images showing FISH result of cell releasing from CIC structures. Layout is same as that of (A). Scale bar: 15 μm. Yellow arrow indicates inner cell that is being released. Blue arrow indicates outer cell. FISH results are presented in the same way as (A). See Movie S7. Related to Figure 6H.

---

Movie S1. The formation of entotic CIC structures

**Entotic Adherence:** MCF10A cell indicated by arrows penetrates into one of its neighboring cells to form CIC structure while being adherent to matrix.

**Entotic Suspension:** Two MCF10A cells indicated by arrows likely first formed CIC structure while being suspended, and then penetrated into one of their neighboring cells to form CIC structure.

**Entotic Division:** Arrow indicated MCF10A cell first underwent cell division with a metaphase of 80 min, and then one of the daughter cells internalized into its neighboring cell to form CIC structure.

**Normal Division:** Normal cell division with a short metaphase of 20 min and two daughter cells adherent to matrix rapidly.

**Movie S2. Entosis is coupled with mitotic cell division.**

Yellow arrows-indicated cell underwent mitotic cell division, followed by internalization of two daughter cells into their neighboring cell (green arrow). The most inner cell died prior to the end of the time lapse.

**Movie S3. Inner cell fates in CIC structures.**

**Death:** Arrow indicated inner cell of a CIC structure died inside another MCF10A cell.

**Unchanged:** Arrow indicated inner cell of a CIC structure stayed alive inside another MCF10A cell to the end of time lapse of 24 h.

**Released:** Arrow indicated inner cell initially resided in the vacuole of a CIC structure, then came out alive as indicated by arrows.

**Movie S4. Catastrophic cell death of mitosis.**

CDC20 depleted mitotic MCF10A cell underwent prolonged metaphase of 480 min and finally died in a catastrophic way.

**Movie S5. Mitosis of different metaphase.**

**M=20 min:** Mitotic MCF10A cell underwent short metaphase of 20 min.

**M=60 min:** Mitotic MCF10A cell with CDC20 depleted underwent prolonged metaphase of 60 min.

**Movie S6. RhoA activity by FRET during one-cell mintosis.**

RhoA activity by FRET in CDC20 depleted MCF10A cell first accumulated during metaphase, then, one of the daughter cells that demonstrated delayed RhoA activity attenuation penetrated into its neighboring cell to form CIC structure, while the daughter cell that rapidly reduced its RhoA activity attached to plate bottom. Left is FRET channel, right is DIC channel.

**Movie S7. Time lapse for FISH.**

**Normal division:** Cell divided with metaphase less than 20 min, two daughter cells attached to plate bottom shortly after division.

**Mintotic division:** Cell divided with one of the two daughter cells internalized to form CIC structure at the end of time lapse, while another daughter cell attached to plate bottom shortly after division.

**Two-cell mintosis:** Cell divided with both of the two daughter cells internalized to form CIC structure at the end of time lapse.

**Inner cell releasing:** The arrow indicated inner cell of a CIC structure started to come out at the end of time lapse.

## Notes

### Competing Interest Statement

The authors have declared no competing interest.

